# The Carbon Dioxide-induced Bioluminescence Increase in *Arachnocampa* Larvae

**DOI:** 10.1101/2020.03.12.989616

**Authors:** Hamish Richard Charlton, David John Merritt

## Abstract

*Arachnocampa* larvae utilise bioluminescence to lure small arthropod prey into their web-like silk snares. The luciferin-luciferase light-producing reaction occurs in a specialised light organ composed of Malpighian tubule cells in association with a tracheal mass. The accepted model for bioluminescence regulation is that light is actively repressed during the non-glowing period and released when glowing through the night. The model is based upon foregoing observations that carbon dioxide (CO_2_) – a commonly-used insect anaesthetic – produces elevated light output in whole, live larvae as well as isolated light organs. Alternative anaesthetics were reported to have a similar light-releasing effect. We set out to test this model in *Arachnocampa flava* larvae by exposing them to a range of anaesthetics and gas mixtures. The anaesthetics isoflurane, ethyl acetate, and diethyl ether did not produce high bioluminescence responses in the same way as CO_2_. Ligation and dissection experiments localised the CO_2_ response to the light organ rather than it being a response to general anaesthesia. Exposure to hypoxia through the introduction of nitrogen gas combined with CO_2_ exposures highlighted that continuity between the longitudinal tracheal trunks and the light organ tracheal mass is necessary for recovery of the CO_2_-induced light response. The physiological basis of the CO_2_-induced bioluminescence increase remains unresolved but is most likely related to access of oxygen to the photocytes. The results suggest that the repression model for bioluminescence control can be rejected. An alternative is proposed based on neural upregulation modulating bioluminescence intensity.

**Summary Statement:** CO_2_ was thought to act as an anaesthetic producing elevated bioluminescence in *Arachnocampa*. Here we show it acts directly on the light organ and does not act as an anaesthetic.

## Introduction

Bioluminescence, the emission of visible light by a living organism as a result of a chemical reaction, occurs in a remarkable diversity of organisms spanning terrestrial and marine environments (Wilson and Hastings, 1998). Among arthropods, bioluminescence has been observed in crustaceans, insects, and myriapods with functions including sexual communication, aposematic signalling, and prey attraction. In all bioluminescent arthropods, light is produced as the result of the luciferin-luciferase chemical reaction (Viviani, 2002). In this reaction, luciferase enzymes catalyse the oxygenation of luciferins to produce electrically excited compounds and photons of visible light (Kahlke and Umbers, 2016).

The best-characterised and most conspicuous terrestrial bioluminescent insects are the fireflies (Order Coleoptera: Family Lampyridae) and the members of the genus *Arachnocampa* (Order Diptera: Family Keroplatidae) (Branham and Wenzel, 2001; Meyer-Rochow, 2007). Among these insects, significant differences in bioluminescence production, utilisation, and regulation have been observed (Lloyd, 1966; Meyer-Rochow and Waldvogel, 1979; Meyer-Rochow, 2007). Adult lampyrid beetles emit light in controlled, periodic, patterned flashes to detect and communicate with potential mates (Copeland and Lloyd, 1983; Lloyd, 1966). Lampyrid larvae release a steady glow, believed to be used aposematically, corelating with distastefulness (Matthysen, 1999). *Arachnocampa* larvae are predators that produce light continuously throughout the night to lure arthropods into web-like silk snares (Broadley and Stringer, 2001; Mills et al., 2016). The light-producing organs in *Arachnocampa* and fireflies are evolutionarily independent and morphologically distinct so bioluminescence production and regulation are expected to differ (Viviani et al., 2002).

The genus *Arachnocampa* is comprised of 9 species endemic to Australia and New Zealand (Baker, 2010; Baker et al., 2008; Meyer-Rochow, 2007). The larvae inhabit cool, dark places including rainforest embankments and the inside of wet caves (Berry et al., 2017; Merritt et al., 2012; Meyer-Rochow, 2007). The lifespan of an adult *Arachnocampa* is very short with adult males living for a maximum of 6 days and adult females living for 2-5 days. The larval state has the longest duration, lasting for many months, during which it utilises its bioluminescence to attract prey (Baker and Merritt, 2003; Merritt and Baker, 2001; Willis et al., 2011). The larvae are relatively immobile and construct snares consisting of mucous-dotted silk lines that hang downward from mucous tubes anchored to a rocky or earthen substrate (Baker and Merritt, 2003; Broadley and Stringer, 2001; Merritt and Baker, 2001; Mills et al., 2016; Willis et al., 2011). The species used in this study, *Arachnocampa flava*, is endemic to south-east Queensland with large epigean populations at Springbrook National Park (Wilson et al., 2004).

### Morphology of the light organ

Light is produced by a posteriorly-located light organ (LO), composed of the modified, large-diameter distal cells of the Malpighian tubules in association with a tracheal mass (Green, 1979; Wheeler and Williams, 1915). The photocytes have a dense cytoplasm with synaptic contacts on the cells of the light organ containing dense-core vesicles that are indicative of neurosecretory regulation (Green, 1979). A single nerve runs from the terminal abdominal ganglion (TAG), separating into neural processes that innervate the LO (Gatenby, 1959; Rigby and Merritt, 2011). The lateral and ventral surfaces of the LO are covered by a mass of tracheoles with interspersed nuclei, taking on a silvery appearance visible through the cuticle (Green, 1979; Rigby and Merritt, 2011). The tracheal layer is closely associated with the photocytes (Green, 1979), suggesting that access to oxygen is a critical factor in bioluminescence output just as it is in fireflies (Ghiradella and Schmidt, 2004); however, the firefly LO is evolutionary derived from a different tissue, believed to be fat body (Amaral et al., 2017).

### Neural control of bioluminescence and effect of anaesthetics

While the regulatory mechanism of firefly bioluminescence is well-known (Ghiradella and Schmidt, 2004; Lloyd, 1966; Timmins et al., 2001; Trimmer et al., 2001), the regulation and production of light by *Arachnocampa* larvae is less well-known. Prior to this study, the prevailing model for bioluminescence regulation in *Arachnocampa* was that bioluminescence is actively repressed when larvae are not glowing such as under daylight or when disturbed, and that the repression is released under darkness (Gatenby, 1959; Rigby and Merritt, 2011). The repression is re-initiated if larvae are exposed to light (Mills et al., 2016). The fact that larvae have a capacity to increase their bioluminescence when stimulated by vibration or the presence of prey in their webs led Mills et al (2015) to propose a two-part system where bioluminescence can also be actively promoted. Evidence for the repression-based model came from ligation and gas exposure experiments. Ligating larvae behind the terminal abdominal ganglion anterior to the LO caused the LO to emit light (Gatenby, 1959) and isolated LOs with neural connections removed emitted low levels of light (Rigby and Merritt, 2011), interpreted as being due to the release of inhibition. The model was reinforced by the response to anaesthetics. In *A. richardsae*, the anaesthetics CO_2_, ether and chloroform caused light release in whole larvae while methanol and ethyl acetate were ineffective (Lee, 1976). Isolated light organs of *A. flava* released low-intensity light when removed from the body and then emitted very bright light when CO_2_ was introduced (Rigby and Merritt, 2011).

From the foregoing studies, one possible explanation for the apparent anaesthetic effect of CO_2_ is that it causes loss of muscle control and that muscle activity could be a requirement for the repression of bioluminescence through modulation of oxygen supply to the LO (Gatenby, 1959; Rigby and Merritt, 2011). It is known that bioluminescence requires oxygen because placing glowing larvae under a vacuum caused them to dim over several minutes (Lee, 1976). In insects, CO_2_ is generally thought to block synaptic transmission at glutamatergic synapses including those at neuromuscular junctions (Badre et al., 2005b); therefore, muscular control could be the key to restricting or activating bioluminescence output. Rigby and Merritt (2011) saw no obvious muscular sphincters associated with the LO or its tracheal supply but acknowledged that this avenue is worthy of further investigation.

Current understanding is that in fireflies neurally-regulated oxygen-gating is involved in the regulation of flash bioluminescence (Timmins et al., 2001). Oxygen transfer to light-producing peroxisomes is interrupted by the mitochondria consuming oxygen between episodes of bioluminescence (Timmins et al., 2001; Trimmer et al., 2001). Flashes occur when uptake of oxygen by mitochondria is repressed by nitric oxide arising from octopamine release. In contrast to adult fireflies, *Arachnocampa* larvae slowly modulate light emission suggesting that the systems will differ (Mills et al., 2016). Further, the two systems have evolved independently from different organ systems. Nevertheless, a common feature is that both fireflies and *Arachnocampa* possess a rich and effective tracheal supply to the LO and both require ATP and oxygen for light release (Kahlke and Umbers, 2016; Mills et al., 2016; Timmins et al., 2001). It appears the access to oxygen is a vital factor for light production in both the coleopteran and dipteran bioluminescence systems.

### Vibration modulation of bioluminescence

*Arachnocampa* larvae brighten substantially when exposed to a vibration stimulus (Mills et al., 2016); vibration of whole larvae in containment produced a 7-10 fold increase in bioluminescence. To incorporate this neurally-based brightening, Mills *et al.* (2016) proposed a two-part regulatory system; (1) a bioluminescence-inhibiting system that prevents bioluminescence when larvae are exposed to daylight or natural light and (2) an acute vibration response, whereby vibration causes an increase in light output followed by a return to pre-stimulus levels. A key aspect of the inhibition hypothesis was the effect of CO_2_ anaesthesia in releasing the proposed inhibitory control, resulting in uncontrolled light output.

Here we explore the physiological and morphological mechanisms of bioluminescence regulation by exposing *A. flava* larvae to a range of anaesthetic, atmospheric, and vibration stimuli and examining the consequent responses. It complements other studies of the *Arachnocampa* luciferin-luciferase system, which are revealing similarities with coleopteran systems – the luciferases belong to the same family of enzymes (Sharpe et al., 2015; Silva et al., 2015) – as well as differences – the *Arachnocampa* luciferin is novel, being different to any described to date (Watkins et al., 2018). Given that bioluminescence is emitted in long bouts and comes under slow neural control, the bioluminescence regulatory mechanisms of *Arachnocampa* are of significant interest when compared to the well-known adult firefly system.

## Materials and methods

### Experimental animals: collection and maintenance

*A. flava* larvae were collected from Springbrook National Park, Queensland, Australia in accordance with a Department of Environment and Science permit (PTU18-001356). Larvae (∼ 2 – 2.5 cm length), probably corresponding to the fourth or fifth instar stages, were collected. Larvae were individually housed in halved, inverted plastic containers (7 cm height × 7 cm diameter) with clay pressed into the upturned base, and fronted with transparent plastic. The containers were kept in glass aquaria filled with ∼1 cm of water and sealed with a glass lid and plastic wrap to ensure high humidity. The aquaria were placed inside a temperature-controlled (24 ± 1°C) room under 12:12 light:dark (L:D) conditions with artificial light produced by a 12V DC white LED lamp controlled by a timer. Larvae were acclimatised to the conditions for a minimum of one week before being utilised in experiments. Larvae that pupated during or immediately following the experiment were not used in analysis. Once per week during the photophase of a non-treatment day (at ∼ 13:00 hours), larvae were fed three CO_2_-anaesthetised *Drosophila melanogaster* adults by securing the *Drosophila* to the silk lines of *A. flava*, taking care to minimise damage to the lines.

### Measuring bioluminescence

Larval bioluminescence was imaged using a SLR camera (Canon EOS 1000D), using identical exposure time, lens and distance between the lens and larva for all experiments. Camera settings were 25 seconds exposure, F5.6, ISO 640. A Pclix XT intervalometer camera controller was used to define intervals between exposures. The application, ImageJ (Version 1.52a), was used to analyse the light output from the individual larvae using the method described by (Mills et al., 2016). Briefly, the greyscale images were imported into ImageJ, then a threshold value (Max: 250, Min: 30) was applied to distinguish the image of the glowing LO region from the background and the intensity of the pixels comprising the glow was summed. As the intensity of isolated pixels is a non-SI unit, the bioluminescence output is referred to in terms of relative light units.

### Light output of live larvae

To investigate the effect of CO_2_, 4 live larvae were pinned to a wax dish, taking care not to damage any internal structures. They were then placed under the camera in a transparent, airtight plastic container (400 ml) with gas ingress and egress connectors and imaging initiated. The pre-treatment bioluminescence output was recorded once per minute for 15 minutes, then larvae were exposed to the treatment gas for 1 minute at a rate of 10 L/minute. The inflowing CO_2_ was humidified by bubbling through water. Imaging continued for 15 minutes after treatment.

### Light output in ligated larvae

To investigate light emission when there was no direct connection between the brain and LO, larvae were ligated with a silk thread at two different locations; (1) behind the head or (2) just anterior to the LO, and the LO-containing region separated by cutting anterior to the ligation. The LO section was then placed on a wax dish (head ligation) or in an excavated glass block under saline (2.6 mmol l^-1^ KCl, 1.8 mmol l^-1^ CaCl2, 150 mmol l^-1^ NaCl and 11 mmol l^-1^ sucrose) (Lozano et al., 2001) and exposed to CO_2_ using the chamber described above with a 15-minute pre-treatment recording period followed by a 1-minute CO_2_ exposure, followed by 15 minutes of recording.

### Light output of isolated light organs

To isolate the LO, a larva was pinned to a wax dish and a cut was made along the dorsal midline to expose the internal organs. The LO, visible as a small cell mass associated with the silvery-white tracheal reflector, was removed from the body and placed in saline. For each application (n = 9), a baseline bioluminescence output was established over 15 minutes. The saline was then replaced with CO_2_-saturated saline and imaged for another 15 minutes. CO_2_ saturation was achieved by bubbling the gas through the saline beforehand via an aquarium aerator stone for 60 seconds. As a control, the isolated LOs of 5 larvae were treated with acidified saline at the same pH as the CO_2_-saturated saline (pH 5.26). As an alternative means of introducing CO_2_, the gas was introduced for 1 minute into the air-space above an excavated block holding isolated LOs (n = 8) under saline.

### Light output in semi-intact dissected larvae

An apparatus was designed to allow introduction of gas to exclusively the head region or the LO-containing region of a partly-dissected larva. Larvae (n = 5 for each treatment) were pinned and dissected along the dorsal midline on a wax dish. A plastic cylinder (5 mm diameter) was then placed over either the LO or the head region containing the brain and an air-tight barrier was formed using petroleum jelly to isolate the cylinder from the rest of the larva. A glass cover-slip was then placed over the top of the cylinder. A gas inlet and outlet port were drilled into the side of the cylinder. This approach allowed sequential directed introduction of CO_2_ into the airspace (a) above the light organ or (b) above the brain.

After dissection and mounting of the cylinder, a larva was imaged for 15 minutes (1 frame per minute) and CO_2_ was introduced into the cylinder air-space for 1 minute (10 L/min). The larva was then imaged for a further 25 minutes and the CO_2_ cylinder was moved to the alternative target area (either LO or brain) not exposed in the initial phase. CO_2_ was introduced at the second location and imaged for 15 minutes.

To apply CO_2_ in solution, larvae (n = 5 for each treatment) were pinned and dissected on a wax dish to reveal the LO and internal organs, as described above. Before any saline was applied, a petroleum jelly barrier was constructed across the mid-body so that a solution applied to either segment would not contact or flow through to the other, then both the anterior and posterior regions were immersed in a drop of saline. The larvae were imaged for 15 minutes then CO_2_-suffused saline was placed on either (a) the LO segment or (b) the brain segment and imaged for a further 15 minutes. Standard saline was used to flush away the CO_2_-suffused saline before imaging for a further 25 minutes. CO_2_-suffused saline was then placed on whichever target site was not exposed in the initial phase of the experiment and imaged for a further 15 minutes.

### Effect of hypoxia on light output

To investigate the effect of hypoxia, the isolated light organs of 3 larvae in saline were exposed to humidified N_2_ in the air-space above the LOs for 1 minute. After 15 minutes, CO_2_ was introduced for 1 minute at a rate of 10 L/minute. In a modified approach, isolated LOs (n = 3) in saline were exposed to N_2_ for 1 minute followed immediately by CO2 for 1 minute, both at a rate of 10 L/minute. After 15 minutes the CO_2_ exposure was repeated. This treatment was also applied to ligated, head-removed sections of larvae (n = 3). Hypoxia recovery was further explored by exposing either isolated LOs (n = 9) or ligated, head-removed sections of larvae (n = 9) to N_2_ for 1 min at a rate of 10 L/minute, followed by a recovery time of either 5, 10, or 15 minutes before being exposed to CO_2_ at the same rate and duration. The imaging set-up and pre-and post-exposure periods were the same as for CO_2_ exposures described above.

### Alternative anaesthetics

To investigate bioluminescence output when the LO was exposed to alternative anaesthetics, isolated LOs in saline were exposed to ethyl acetate or diethyl ether (n = 4, for each treatment). Approximately 50 ml of either ethyl acetate or diethyl ether was placed in a 500 ml glass conical flask and vapour allowed to accumulate for 15 minutes. To expose the LO, air was pumped through this bottle into the LO-chamber for 1 minute at a rate of 10 L/minute with a pre- and post-exposure imaging period of 15 min. An isoflurane vaporiser (Model V300PS) was used to expose the larvae (n = 4) to 4% isoflurane at the same rate and duration as the other anaesthetics.

### Vibration response

Larvae were placed in a humidified aquarium before the onset of darkness. Two 10 mm diameter vibrating AD1201 haptic motors (180-200 Hz) were attached to the sides of the aquarium with duct tape and operated using a programmed Raspberry Pi 3 (Model B V1.2). Light output of the larvae was recorded at 1-minute intervals through the night using a SLR camera (Canon EOS 1000D) under time-lapse control, viewing the larvae from below the aquarium. The vibrating haptic motors were activated in 15 sec pulses at a range of inter-pulse intervals throughout the scotophase (Table 1). Treatment nights were interspersed between control nights with no vibration stimuli, using the same larvae. Light intensity of individual larvae was determined using ImageJ as described above. In this series of experiments the light units are not comparable to those from all other experiments due to the different location of the camera. The light output was displayed as the mean intensity of the larvae in the aquarium. The same larvae were used in multiple treatments; however, data from larvae that pupated during or within three days of an experiment were removed.

### Statistical analysis

Data from physiological experiments were analysed using a paired t-test, comparing the total mean light output for 15 minutes pre- and post-exposure to stimuli. Further, a paired t-test was performed to compare light output in paired trials on the same larvae, as this test compares two different methods of treatments, where the treatment is applied to the same subject. The test determined if the two methods of treatment produced results with significant difference.

## Results

### CO_2_ response

The bioluminescence output of live larvae (n = 4) increased when exposed to CO_2_ in the air-space (Fig. 1a). Very low levels of light were emitted by the larvae before treatment, but this output did not register on the scale that shows the subsequent response to CO_2_. On exposure, the larval light intensity increased over the next 10 minutes and started to decline after reaching a mean peak light intensity of 6.1 × 10^6^ units.

**Figure 1:**
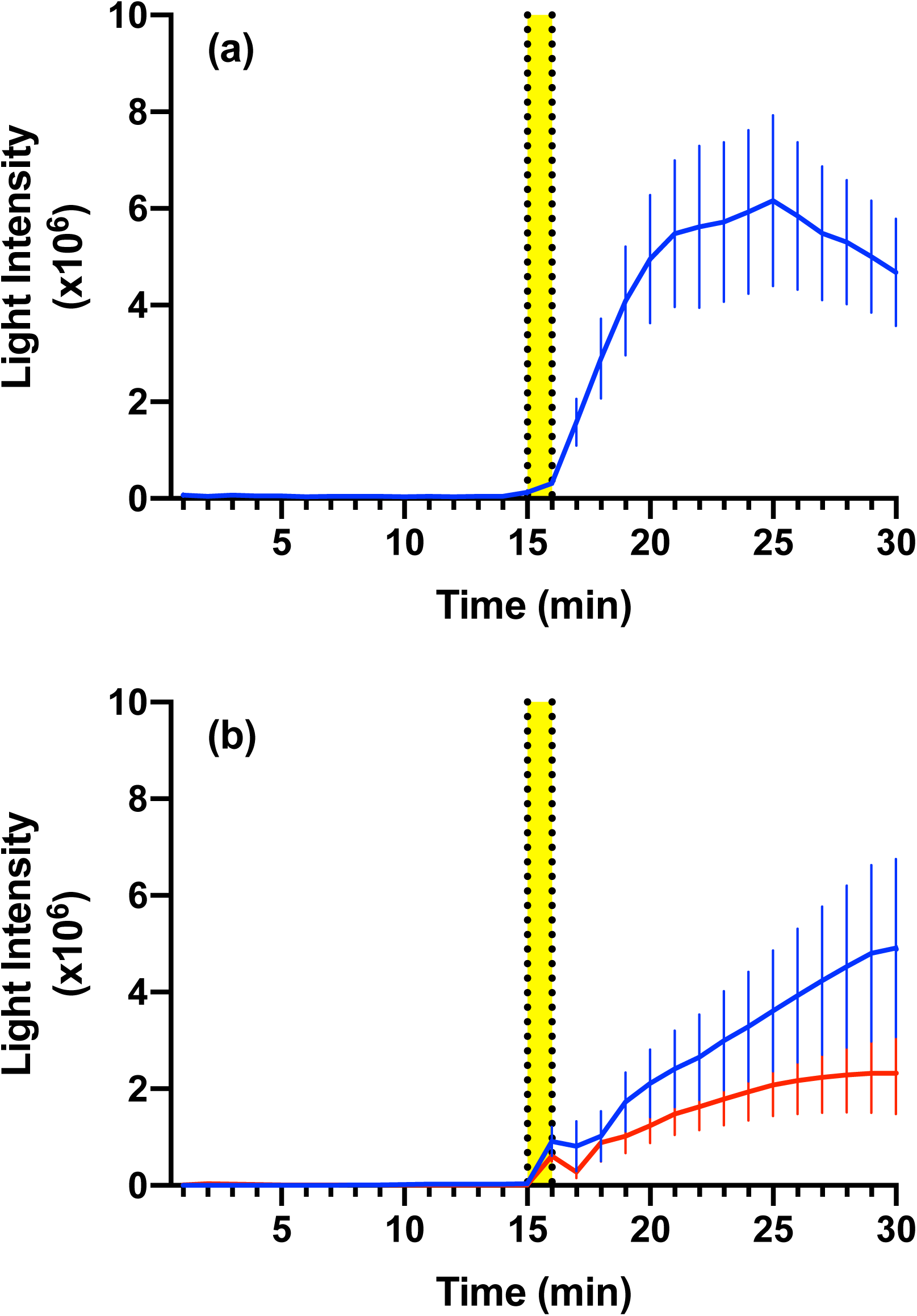
The bioluminescence output of **(a)** live *A. flava* larvae (n = 4, mean ± s.e.m.), and **(b)** ligated, head-removed larvae (n = 10, blue) and ligated LO-isolated larvae (n = 11, red) (means ± s.e.m.) following exposure to CO_2_ in the air-space. Yellow bar indicates period of gas exposure.

Ligated, head-removed larvae and ligated, LO-isolated larvae (n = 10, 11, respectively) displayed increases in bioluminescence intensity when exposed to CO_2_ in the air-space (Fig. 1b). Both treatments showed an initial increase in bioluminescence followed by a brief decrease and then a steady increase in bioluminescence, culminating at a mean peak light intensity of approximately 4.9 × 10^6^ units (ligated, head-removed) and 2.5 × 10^6^ units (ligated, LO-isolated). In both, the bioluminescence level remained elevated after the 15-minute recording period.

When light organs were removed by dissection, they were isolated from the larval body and all contact with nerves and main tracheal trunks removed. After this treatment, the light organs released light at relatively low levels through the pre-exposure period (mean of 3.9 × 10^5^ units). Exposure to CO_2_-saturated saline (n = 9) or CO_2_ in the air-space (n = 8) produced an immediate increase in bioluminescence intensity within 1–2 minutes (Fig. 2). The CO_2_-saturated saline treatment produced a peak bioluminescence level of 7.1 × 10^6^ units, that exponentially declined over the next 15 minutes but remained above the pre-stimulus levels of bioluminescence. The CO_2_-air-space treatment showed a lower peak of 5.5 × 10^6^ units and returned to pre-stimulus bioluminescence levels after approximately 7 minutes.

**Figure 2:**
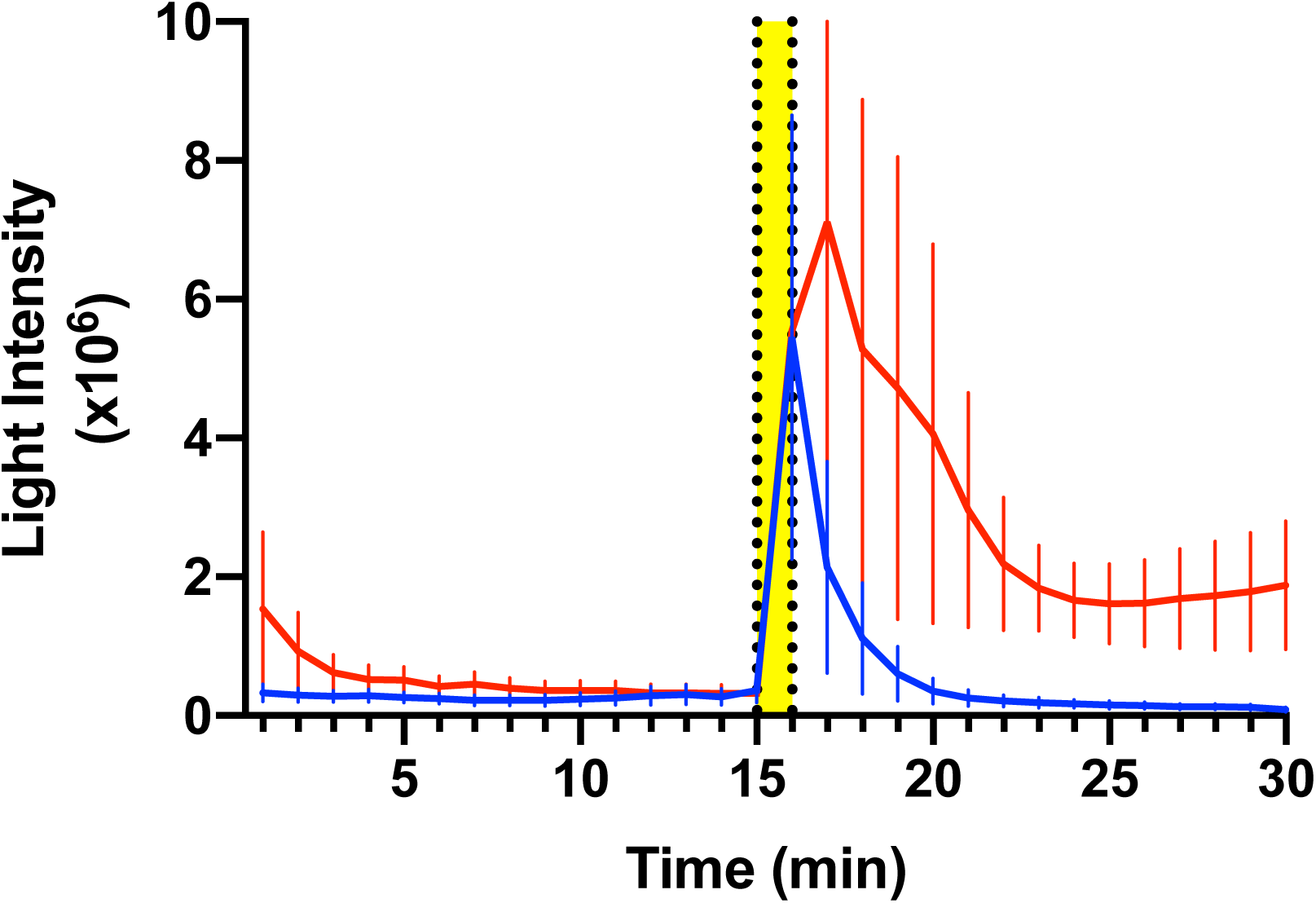
The bioluminescence outputs of isolated *A. flava* LOs following exposure to CO_2_-saturated saline (n = 9, red) and the introduction of CO_2_ into the air-space (n = 8, blue) (means ± s.e.m.). Yellow bar indicates CO_2_-diffused saline application or introduction of CO_2_ into the airspace.

### Localisation of the CO_2_ effect

The high light emission response to CO_2_ was localised by sequentially exposing the LO and the brain to CO_2_ using two different approaches (1) applying CO_2_-saturated saline to either region or (2) exposing the air-space above either region. When CO_2_-saturated saline was applied to the LO first (n = 5) (Fig. 3a) bioluminescence production increased greatly and no obvious increase occurred when the treatment was applied to the brain. Similarly, when the CO_2_-saturated saline was applied to the brain first and then the LO (n = 5), bioluminescence intensity increased when the treatment was directed to the LO. The sequential application of CO_2_ to the air-space above the brain and the LO (n = 5 for each treatment) (Fig. 3b) produced similar results: exposure to the brain region produced no bioluminescence response while exposure to the LO produced a large response. Overall, the bioluminescence responses recorded through exposure to CO_2_-saturated saline (Fig. 3a) produced less light than those exposed to CO_2_ in the air-space (Fig. 3b). Air-space exposure produced a shorter-lasting light output than saline exposure.

**Figure 3:**
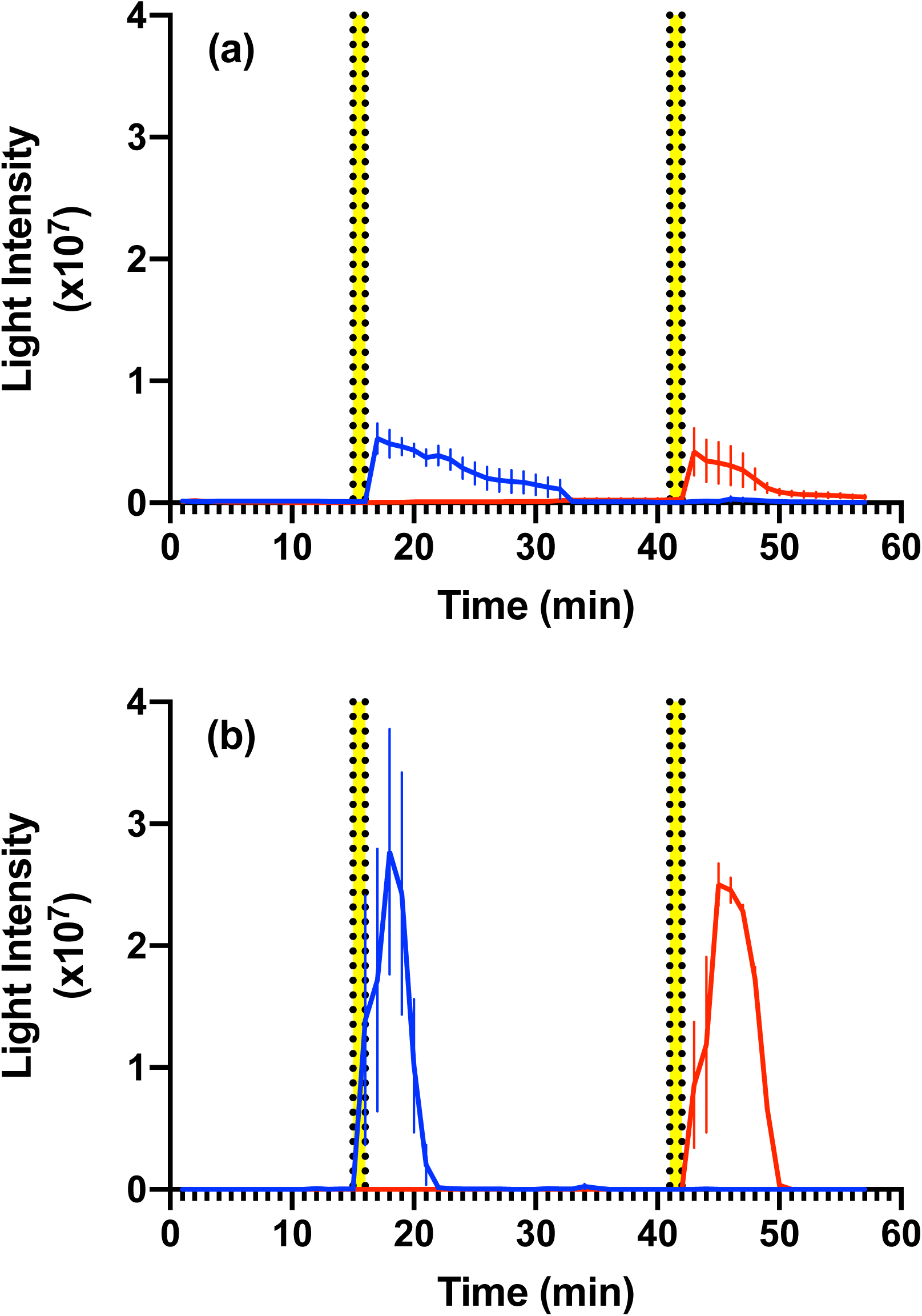
The bioluminescence outputs of semi-intact *A. flava* larvae when exposed to **(a)** CO_2_-saturated saline (yellow bars) in the order **(1)** the LO followed by the brain (n = 5, blue) or **(2)** the brain followed by the LO (n = 5, red) (means ± s.e.m.). **(b)** Bioluminescence outputs following air-space CO_2_ exposure (yellow bars) depicted as the same colour codes as **(a)** (n = 5, for each treatment order) (means ± s.e.m.).

To determine if the application of CO_2_ was indirectly triggering acute bioluminescence response by altering either the internal pH of *A. flava* or the pH of the saline, an acidified saline of pH 5.26, identical to that of CO_2_-saturated saline, was applied to isolated LOs (n = 4). No bioluminescence responses were recorded after exposure (data not shown).

### Combining hypoxia and CO_2_ exposure

To determine the effect of hypoxia on bioluminescence production, isolated LOs (n = 3) were exposed to N_2_ for 1 minute, immediately followed by the introduction of CO_2_ for the same period. No bioluminescence responses were detected (data not shown). When a 15-minute recovery period was allowed, and a second CO_2_ treatment was applied, the isolated LOs (n = 3) produced no bioluminescence responses (data not shown). However, when this same treatment series was applied to ligated, head-removed larvae (n = 3) (Fig. 4), a low response was elicited after the first CO_2_ exposure and a much higher bioluminescence level was recorded following the second CO_2_ exposure. As an additional control, introduction of N_2_ into the air-space above isolated LOs produced no bioluminescence (n = 3, data not shown).

**Figure 4:**
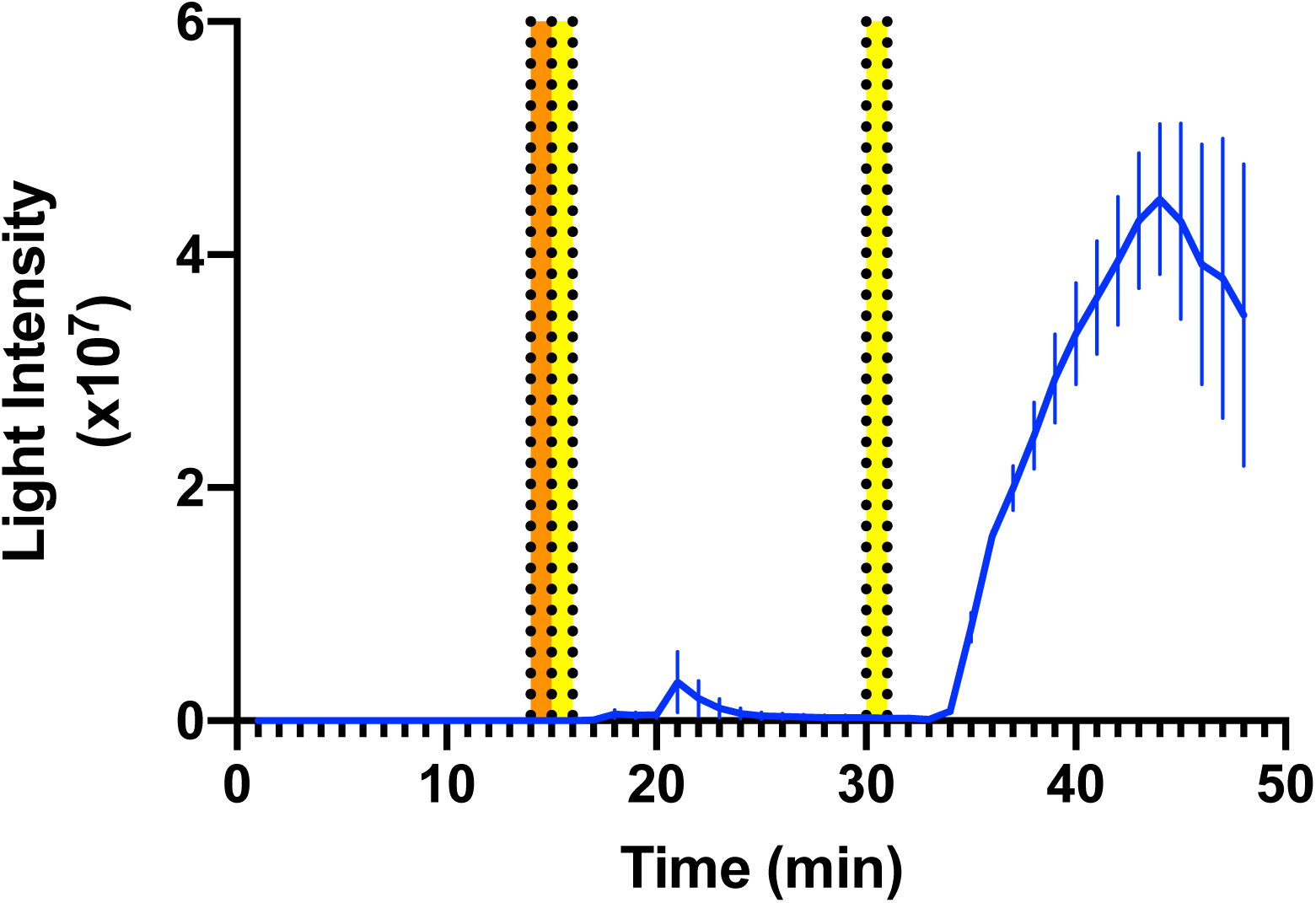
The bioluminescence output of ligated, head-removed *A. flava* larvae (n = 3) in response to N_2_ exposure (orange bar) immediately followed by CO_2_-exposure (yellow bar). After a 15-minute recovery period, the LOs were again exposed to CO_2_ (yellow bar).

To investigate the ability of ligated, head-removed larvae to recover from hypoxia, the segments were exposed to N_2_ for 1 minute, followed by a recovery period of 5, 10 or 15 minutes (n=3 for each treatment) before exposure to CO_2_ for 1 minute. In all cases, light was released on exposure to the CO_2_ pulse with the maximum mean light output increasing with recovery time (Figure 5). When the identical treatments were applied to isolated LOs, no light was emitted after the recovery periods.

**Figure 5:**
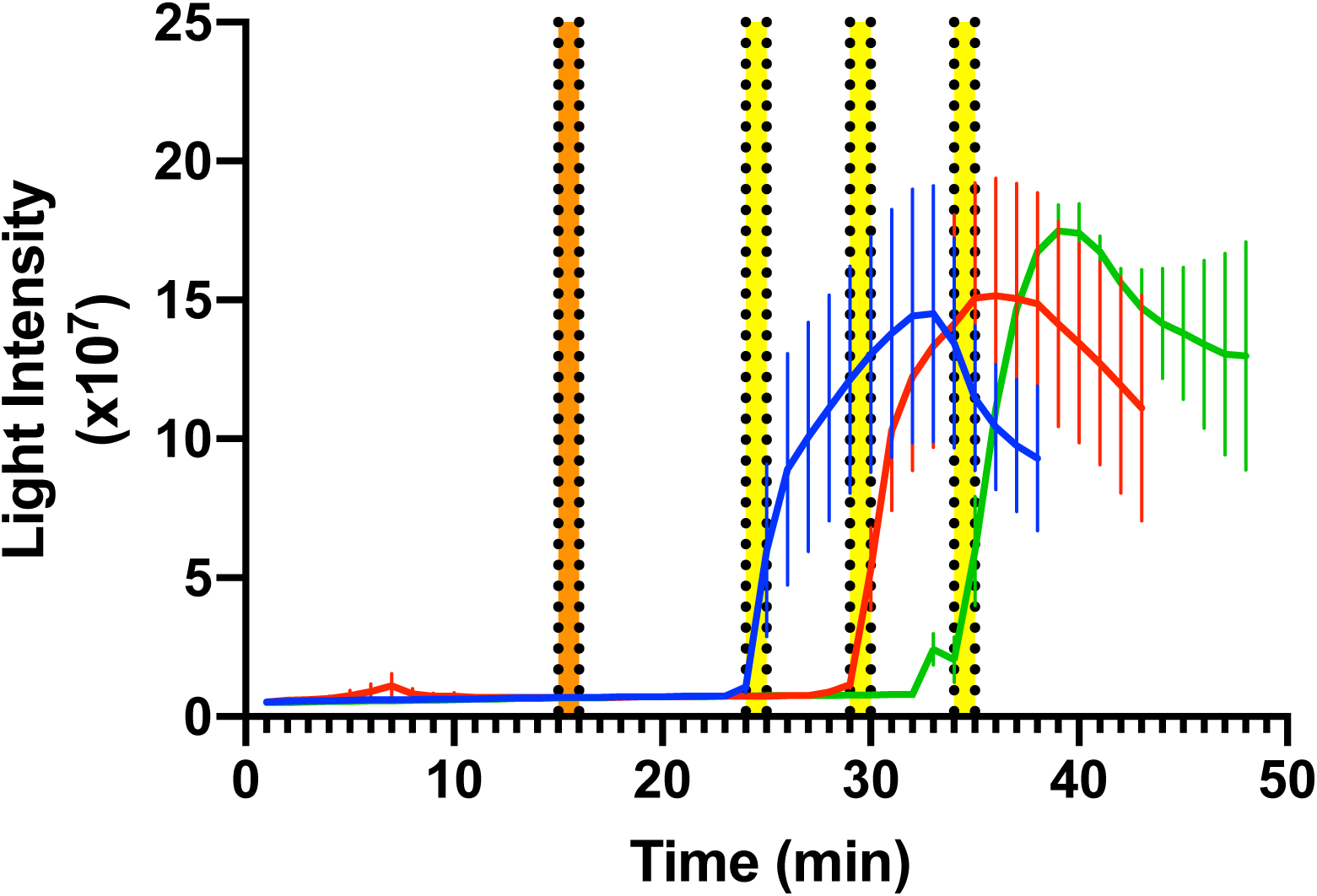
The bioluminescence output of ligated, head-removed *A. flava* larvae (n = 3) in response to N_2_ exposure (orange bar) followed by CO_2_-exposure after 5 (n = 3, blue), 10 (n = 3, red), or 15 (n = 3, green) minutes (yellow bars).

### Alternative anaesthetics

To determine if the CO_2_ response was due to an anaesthetic effect, the alternative anaesthetics isoflurane, ethyl-acetate, and diethyl-ether were applied to isolated LOs via air-space treatment (n = 4 for each treatment) (Fig. 6). None of the treatments produced bioluminescence responses at levels comparable to that induced by CO_2._ A small increase was noted in the ethyl acetate and diethyl ether treatments; however, it did not precisely coincide with exposure.

**Figure 6:**
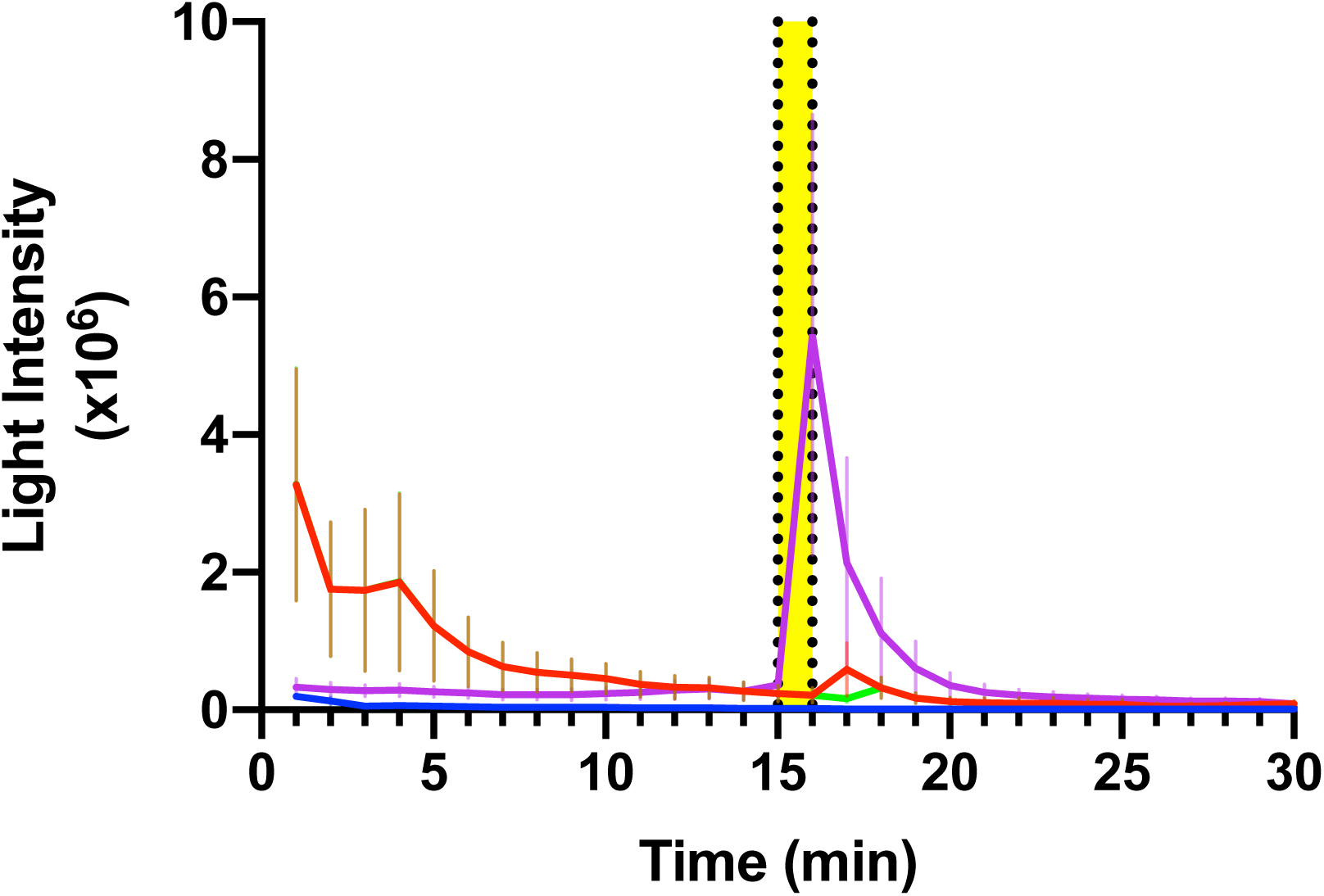
The bioluminescence response of isolated *A. flava* LOs (n = 4 for each treatment) (means ± s.e.m.) following exposure to ethyl-acetate (red), diethyl-ether (green), Isoflurane (blue), and to CO_2_ into the airspace (n = 8) (purple).

### Vibration

Undisturbed larvae began bioluminescence shortly after the onset of darkness and reached peak light intensity approximately 1 hour into the 12-hour scotophase (Fig. 7). Following the peak, the bioluminescence intensity steadily declined over the next 11 hours and did not completely cease until lights-on.

**Figure 7:**
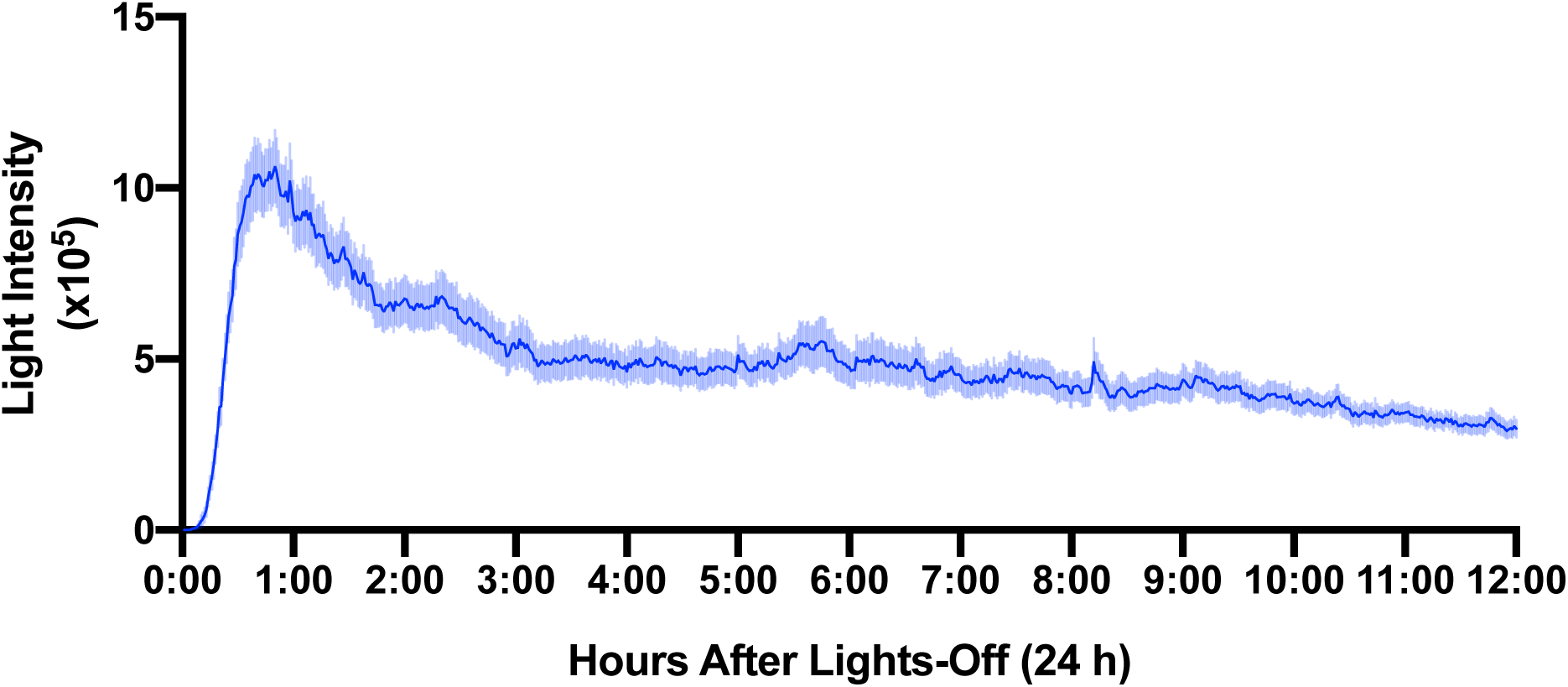
The generalised bioluminescence output (mean ± s.e.m.) derived from 108 *A. flava* larvae recorded in laboratory conditions under an artificial light cycle (lights-off at 0:00 and on at 12:00).

Exposure of larvae to vibration produced significant increases in bioluminescence output compared with the controls recorded during the preceding scotophase without vibration (Paired *t*-toest, *P*<0.001) (Fig. 8). Larvae exposed to one minute of vibration at frequencies of 1 pulse/hour, 1 pulse/30 minutes, and 1 pulse/15 minutes showed acutely elevated light responses to vibration stimulus (Fig. 8a–c). The bioluminescence outputs peaked following vibration and then exponentially returned to approach the pre-stimulus levels. It is apparent that the average amplitude of the acute vibration response decreased as the interval between pulses shortened. Acute responses followed by declines were not seen when treatments occurred less than 15 minutes apart (Fig. 8d–f). Rather, larvae displayed persistent, elevated levels of bioluminescence without the response-decline seen when treatments were more widely spaced. Larvae vibrated at 1 pulse/minute recorded the greatest bioluminescence increase.

**Figure 8:**
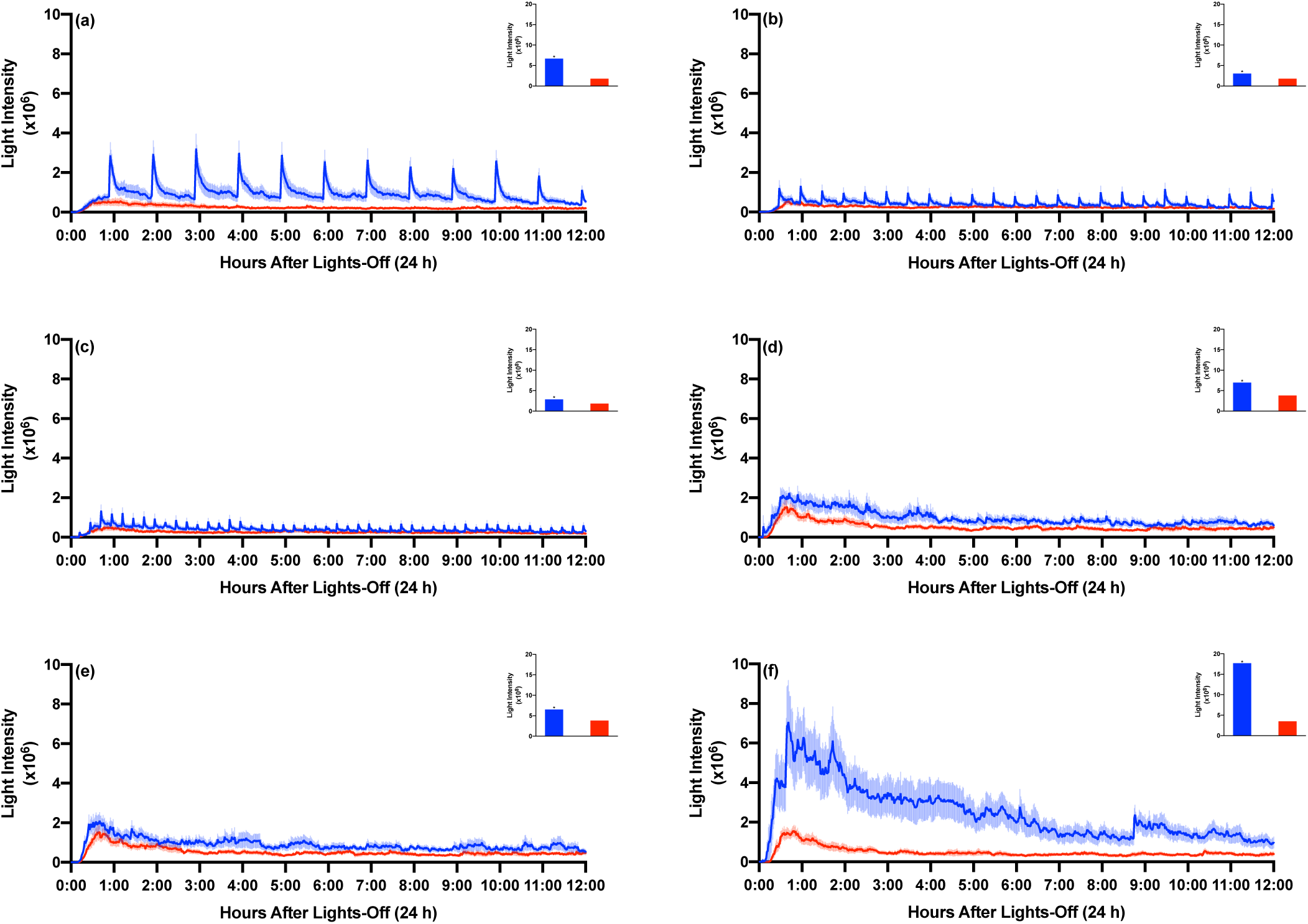
Bioluminescence output (mean ± s.e.m.) of live *A. flava* larvae in response to a 15-second (180-200 Hz) vibration at a rate of: **(a)** 1 pulse/hour (n=17), **(b)** 1 pulse/30 min (n=17), **(c)** 1 pulse/15 min (n=17), **(d)** 1 pulse/12 min (n=7), **(e)** 1 pulse/10 min (n=8), and **(f)** 1 pulse/min (n=9). The bioluminescence output of the same larvae recorded the during prior night’s scotophase is shown in red. The insets show the total bioluminescence output of larvae through the treatment night (blue) in comparison to the total bioluminescence released in the previous control during night (red) (* *P* < 0.001).

When larvae (n = 9) were exposed to vibration pulses at 1 pulse/5 minutes for 3 hr, the bioluminescence level was elevated and responses to the individual pulses were not apparent (Fig. 9). When vibration ceased between 3:00 and 6:55 hours after lights-off, bioluminescence returned to a lower baseline level and approached that recorded in the control treatment. When the pulse series was re-initiated, the larval light output immediately elevated and remained higher than in the rest period.

**Figure 9:**
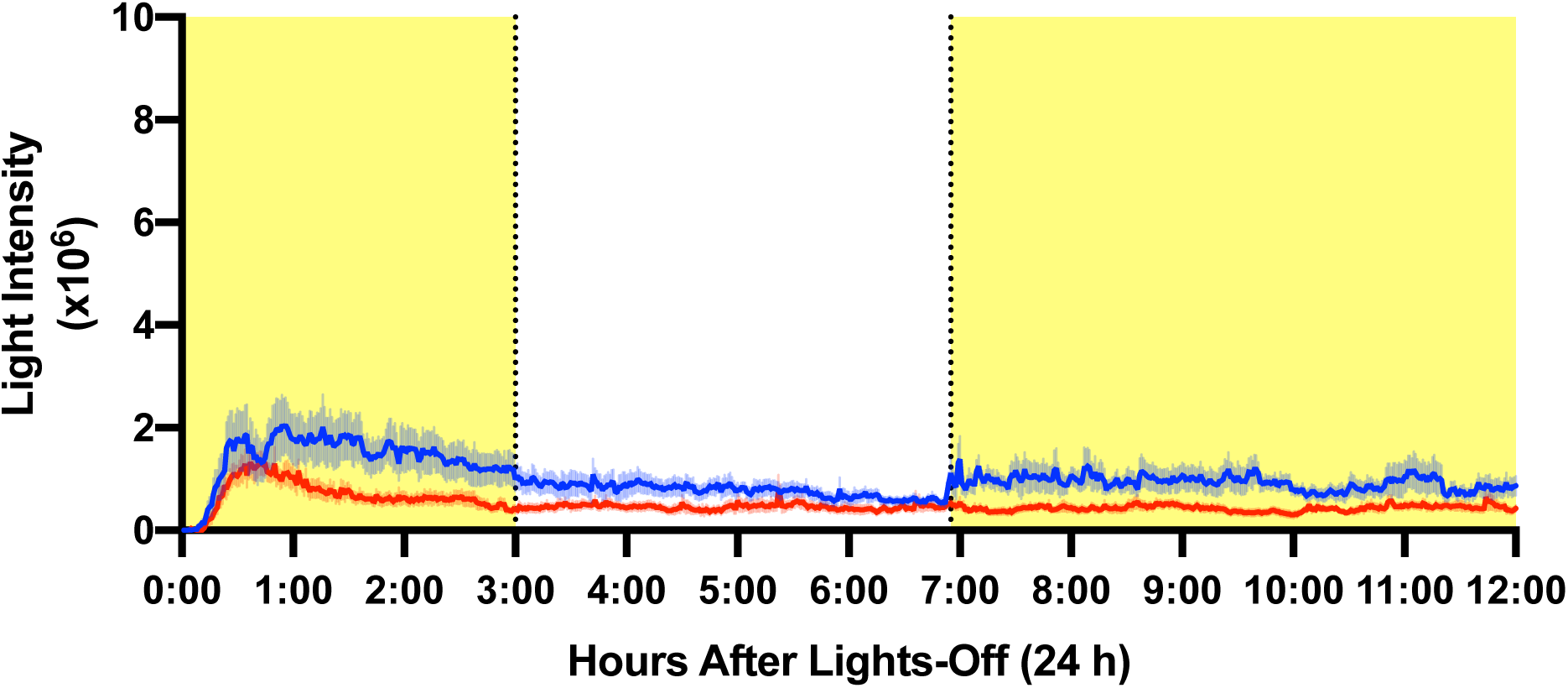
Bioluminescence output (mean ± s.e.m.) of 9 live *A. flava* larvae in response to a 15-second 180-200 Hz vibration at a rate of 1 pulse/5 minutes between 0:00-3:00 and 6:55-12:00 (yellow). Vibration responses were compared to the bioluminescence output of an untreated control (red) recorded the previous night.

## Discussion

### Elevated Bioluminescence in Response to CO_2_

At the initiation of this work, it was believed that CO_2_ causes a major release of light in *Arachnocampa* through its anaesthetic effect (Lee, 1976; Rigby and Merritt, 2011). In insects, CO_2_ is a widely used anaesthetic that is believed to block synaptic transmission at the neuromuscular junction by influencing glutamate sensitivity (Badre et al., 2005). Diethyl ether is a common anaesthetic used for many years on both vertebrates and invertebrates (Whalen et al., 2005). The halogenated ether, isoflurane, is also widely used, including on insects, where it acts upon voltage-gated K^+^ channels (Barber et al., 2012; McMillan et al., 2017). Ethyl acetate is another insect anaesthetic (Loru et al., 2010); however, its mode of action is unknown. Here, we tested the effect of these anaesthetics on different preparations of the light organ and found low to zero response to any other than CO_2_, casting doubt on the concept of anaesthesia releasing neural repression of light output. Isoflurane, ethyl acetate, and diethyl ether produced no bioluminescence responses in isolated light organs that approach the CO_2_-induced response magnitude. A small increase was noted in the ethyl acetate and diethyl ether treatments; however, it did not precisely coincide with exposure. Lee’s (1976) experiments were performed on an unstated and possibly small sample size and responses to the different anaesthetics were not quantified, so it is possible that light levels were not strictly compared among treatments. It has been noted since that the light released upon separation of the LO from the body was very dim compared to that released under exposure of the LO to CO_2_ (Rigby and Merritt, 2011). The present study confirms that observation, that CO_2_ produces intense light production.

To explore the location of the CO_2_ effect, either the brain or the LO of whole, semi-dissected larvae (the body wall was opened up and pinned out under saline to reveal the internal organs) was exposed to either CO_2_-saturated saline or CO_2_ in the airspace. Using either method, bioluminescence responses were seen only when CO_2_ was directed to the LO, indicating that its effect is initiated at the LO and that exposure of the brain has no direct effect on bioluminescence. The outcome also supports the above: that the CO_2_-induced elevation of light is not due to a general anaesthetic effect. The findings cast doubt on the validity of the current models where neural signals from the brain repress bioluminescence and where CO_2_ acts as a general anaesthetic.

How then does CO_2_ have such a strong effect on bioluminescence? A supply of air to the larva is essential for light emission as shown by dimming of larvae under vacuum, and a return when air was readmitted (Lee, 1976). Previous studies have highlighted the intimate relationship between the LO and the tracheal mass (Green, 1979), indicating that the availability of oxygen is necessary for the oxidative bioluminescence reaction, just as it is in firefly larvae (Carlson, 1965; Hastings and Buck, 1956) and adults (Buck, 1948; Timmins et al., 2001). Here, we confirmed that hypoxia limits the CO_2_-induced brightening. When CO_2_ was introduced immediately following N_2_ exposure or after recovery periods of 5 – 15 minutes, the isolated light organs showed no bioluminescence. When CO_2_ was applied to ligated, head-removed larvae immediately after N_2_, a low light output was recorded and a second CO_2_ exposure 15 minutes later produced a significantly higher light output, indicating that the ligated section is capable of recovery from hypoxia. Using another approach, ligated head-removed larvae exposed to N_2_ followed by CO_2_ after a spaced delay (5 – 15 minutes) showed a recovery the intense CO_2_ response, with signs of a graded increase in magnitude with increased recovery time.

We consider it most likely that recovery is due to the LO tracheal mass maintaining a connection with the main longitudinal tracheal trunks in ligated larvae, whereas that connection had been removed in isolated light organs. *Arachnocampa* larvae are apneustic (Ganguly, 1959), i.e. they have no spiracles, although the longitudinal tracheal trunks are air-filled. This respiratory strategy is common in aquatic or endoparasitic insect larvae. Gas exchange is cutaneous—across the cuticle—which is probably aided in *Arachnocampa* by the fact that larvae have thin cuticle and dwell inside a mucous tube. In apneustic insects, air-filling is attributed to the cells lining the trachea having the ability to withdraw fluid from the tracheal lumen and replace it with air (Buck and Keister, 1955). In *Arachnocampa* larvae, the longitudinal tracheal trunks run directly into the tracheal mass associated with the photocytes (Green, 1979). We propose that the epithelial cells of the main paired longitudinal tracheal trunks modulate the oxygen content of the lumen through active transport. The recovery seen in ligated larvae is attributed to reoxygenation of the tracheal lumen during the recovery period and suggests that available oxygen is consumed during the CO_2_-induced bioluminescence.

There have been no reports of similar acute bioluminescence outputs in larval or adult fireflies upon CO_2_ exposure (Buck, 1948) so *Arachnocampa’s* response appears to be unique to its regulatory system. CO_2_ is known to be a product of light production in the ATP-dependent light production system of fireflies (Plant et al., 1968), which is likely to also apply to *Arachnocampa* because of the related luciferase system (Lee, 1976). It could be that light production in the photocytes is modulated by both CO_2_ and O_2_ levels and that the light regulation mechanism is reliant on a balance between both gases. It is possible that high PCO_2_ triggers a runaway reaction due to the supra-metabolic levels of CO_2_ producing maximum light production in the LO cells. Such high and persistent light levels were also seen when larvae were fed prey items dosed with either phentolamine or octopamine, involving biogenic amines in bioluminescence regulation (Rigby and Merritt, 2011) but how neurotransmitters interact with gas regulation remains unknown.

### Vibration response

Larvae are capable of upregulating the level of bioluminescence is situations where prey are detected in their web, during aggressive interactions with other larvae (Mills et al., 2016) and at the onset of rainfall (Merritt and Patterson, 2018). These responses are believed to be mediated through detection of vibration and this stimulus-response system is readily manipulated in the laboratory (Mills et al., 2016). When larvae were glowing at an undisturbed level during the night, a vibration stimulus produced an acute increase in light output to about x times the base level that slowly returned to pre-stimulus levels (Mills et al., 2016). The up-regulation is likely mediated through the nervous system; however, it is not known whether the elevated light emission is attributable to an increased oxygen supply to the photocytes or to the availability of the light-producing reaction involving luciferin, luciferase or ATP. Here we examined whether persistent stimulation would produce habituation or effector fatigue. First, we tested whether light emission decreased with repeated vibration exposures. When larvae were exposed to 1 pulse/15 minutes or less frequently, an acute response was recorded. When pulses were more closely spaced, light output remained persistently elevated above the control level. This appears to be due to a general elevated excitation level because a 3-h gap in stimulus exposures resulted in a gradual decline in light output, approaching the control level, followed by a return to a higher level when stimuli were reinstated. We conclude that habituation does not occur through a 12 h scotophase because the level of response to spaced pulses was consistent. Second, evidence of effector fatigue was seen: the more widely spaced pulses produced consistently higher peak responses. This could be due to reduced availability of light-producing metabolites as the stimuli become more closely spaced. While the experimental observations presented here do not shed light on the mechanism by which vibration elevates light output, the vibration response will be a useful readout for quantifying the effect of gas mixtures on the bioluminescence system.

### Bioluminescence regulatory mechanism

Our findings suggest that the model of light production being suppressed under neural control when it is in the off state should be rejected due to (1) the lack of a bioluminescence response when anaesthetics other than CO_2_ were applied to the LO, and (2) the fact that CO_2_ directly affects the light organ. The vibration responses mentioned above are indicative of an active elevation of bioluminescence in response to a neural stimulus mediated by the sensory system, so a simpler control model is that excitatory neural signals initiate bioluminescence, maintain it at a steady state, and mediate the vibration-induced startle reflex. Octopamine is the prime candidate as the neurotransmitter (Rigby and Merritt). The time-course of light elevation and reduction suggests that the neuronal control acts through a second mechanism acting over a longer time-course with the most likely being the modulation of oxygen access to the photocytes. One possibility is a sphincter-based control system that modulates airflow between the longitudinal trachea and the tracheal mass, as mentioned by others (Gatenby and Ganguly, 1956; Rigby and Merritt, 2011) but not yet thoroughly investigated. Another is modulation of oxygen transfer across the junction between the tracheal mass and the photocytes, perhaps akin to the indirect way oxygen transfer between tracheal end cells and photocytes is modulated in adult fireflies (Timmins et al., 2001; Trimmer et al., 2001). At the ultrastructural level, the tracheolar units are very closely opposed to the basal lamina of the photocyte cells with many deep infoldings of the membrane (Green, 1979), all consistent with a functional association between the photocytes and the tracheal supply. There is no barrier of mitochondria between the photocyte cytoplasm and the air supply as seen in firefly LOs—the large mitochondria in *Arachnocampa* photocytes are distributed throughout the cytoplasm (Green, 1979)— however the firefly arrangement appears to be specifically adapted to flash control. Further experiments exposing larvae to combinations of gas mixtures and excitatory/inhibitory stimuli should reveal more details of light regulation in *Arachnocampa*.

## Abbreviations

CO_2_: Carbon dioxide
N_2_: Nitrogen
LD: Light:dark
SLR: Single lens reflex
TAG: Terminal abdominal ganglion

## Acknowledgements

We would like to thank Dr. Simone Blomberg for statistical advice, Professor Bruno Van Swinderen and Michael Troup for use of isoflurane vaporiser equipment, and members of the Asgari Lab for help with pH measurement. We would like to thank Rory Charlton for his invaluable assistance in designing and constructing the vibration apparatus.

